# A new standard for crustacean genomes: the highly contiguous, annotated genome assembly of the clam shrimp Eulimnadia texana reveals HOX gene order and identifies the sex chromosome

**DOI:** 10.1101/222869

**Authors:** James G. Baldwin-Brown, Stephen C. Weeks, Anthony D. Long

## Abstract

Vernal pool clam shrimp (*Eulimnadia texana*) are a promising model system due to their ease of lab culture, short generation time, modest sized genome, a somewhat rare stable androdioecious sex determination system, and a requirement to reproduce via desiccated diapaused eggs. We generated a highly contiguous genome assembly using 46X of PacBio long read data and 216X of Illumina short reads, and annotated using Illumina RNAseq obtained from adult males or hermaphrodites. 85% of the 120Mb genome is contained in the largest 8 contigs, the smallest of which is 4.6Mb. The assembly contains 98% of transcripts predicted via RNAseq. This assembly is qualitatively different from scaffolded Illumina assemblies: it is produced from long reads that contain sequence data along their entire length, and is thus gap free. The contiguity of the assembly allows us to order the HOX genes within the genome, identifying two loci that contain HOX gene orthologs, and which approximately maintain the order observed in other arthropods. We identified a partial duplication of the Antennapedia gene adjacent to the few genes homologous to the Bithorax locus. Because the sex chromosome of an androdioecious species is of special interest, we used existing allozyme and microsatellite markers to identify the *E. texana* sex chromosome, and find that it comprises nearly half of the genome of this species. Linkage patterns indicate that recombination is extremely rare and perhaps absent in hermaphrodites, and as a result the location of the sex determining locus will be difficult to refine using recombination mapping.

## Introduction

The clam shrimp *Eulimnadia texana* has, along with other vernal pool shrimp, been noted for its unique sex determining system (Sassaman and Weeks 1993), its rare (in Metazoa) requirement to reproduce via desiccated diapaused eggs (Sassaman and Weeks 1993), and its unique habitat. This androdioecious (Sassaman and Weeks 1993) species has three common arrangements of sex alleles (Sassaman and Weeks 1993) or “proto-sex chromosomes” (Weeks et al., 2010). Males are always homozygous for the “Z” male allele, while hermaphrodites may be “ZW” or “WW”, with WW hermaphrodites only capable of producing hermaphrodite offspring. Much effort (S. C. Weeks et al. 2010) has gone into attempting to identify the *E. texana* sex locus because of this unique arrangement; this, coupled with the fact that close relatives of the species have ordinary male-female sexual dimorphism (S. C. Weeks et al. 2009), makes the Eulimnadia clade, and *E. texana* in particular, an excellent study system for understanding the genetic changes that underlie changes in sex determination. The fact that, unlike in most animals, both the “Z” and “W” sex determination alleles are capable of being homozygous is interesting as a comparator for testing the hypothesis that the lack of recombination in “Y” and “W” alleles drives degradation of sex chromosomes. The ability of eggs to remain in diapause for years at a time (Brendonck 1996) is especially valuable to geneticists because very few macroscopic animals exist in which populations can be archived for long periods without changes occurring in the genetics of the population (genetic drift, loss of linkage disequilibrium, etc.). Furthermore, clam shrimp live in desert vernal pools; naturally limited migration from pool to pool makes them well suited to the study of populations evolving in relative genetic isolation.

Genome assembly of non-model organisms was financially unrealistic until the advent of high-throughput next generation sequencing. Unfortunately, next generation sequencing methods such as Illumina are limited to short read sequencing, which is not ideal for genome assembly; assemblies produced using Illumina-type short read data tend to have low contiguity (Treangen and Salzberg 2011). This problem can be overcome by using PacBio (Eid et al. 2009), Oxford Nanopore (Laver et al. 2015), or other long read sequencing technologies to supplement or replace Illumina sequencing. A hybrid approach to sequencing and assembly using both short and long reads has been shown to produce highly contiguous assemblies in *Drosophila-sized* genomes (Chakraborty et al. 2016). Genome annotation of *de novo* assemblies is routinely performed using RNAseq data (Z. Wang, Gerstein, and Snyder 2009), and tools for that purpose are already available (Stanke and Waack 2003; Grabherr et al. 2011).

Here, we lay out our attempt to extend genetic research on *E. texana* into the world of whole genome sequence analysis using the latest genomics techniques. We used a combination of short read Illumina (R. Shen et al. 2005) and long read PacBio (Eid et al. 2009) sequencing to generate a high quality draft genome assembly and performed an annotation of genes using RNAseq (Z. Wang, Gerstein, and Snyder 2009). We generated a genome assembly for a WW hermaphrodite clam shrimp strain consisting of 112 contigs totaling 120Mb in length with a contig N50 of 18Mb. Using RNAseq data we annotate 17,667 genes, of which ~99% of hermaphrodite transcripts are placed into our assembly. This assembly is the most contiguous assembly of a crustacean genome of which we are aware. By comparison, *Daphnia pulex* has a scaffold N50 of 494kb (Ye et al. 2017).

## Methods

### Shrimp collection and rearing

Clam shrimp (Fig. 1) were sampled from Arizona and New Mexico as previously described (S C Weeks and Zucker 1999). We reared the clam shrimp in the laboratory until day 10 of their life cycles, then extracted DNA and RNA from them. Clam shrimp populations were reared in 50X30X8 cm disposable aluminum foil catering trays (Catering Essentials, full size steam table pan). In each pan, we mixed 500mL of soil with 6L of water purified via reverse osmosis. 0.3 grams of aquarium salt (API aquarium salt, Mars Fishcare North America, Inc.) were added to each tray to ensure that necessary nutrients were available to the shrimp. Trays were checked daily for non-clam shrimp, especially the carnivorous *Triops longicaudatus*, and all non-clam shrimp were immediately removed from trays. We identified the following non-clam shrimp: *Triops longicaudatus, Daphnia pulex*, and an unknown species of *Anostraca* fairy shrimp. An inbred population of clam shrimp, here referred to by its numerical title JT4(4)5, was derived from the JT4 wild population and used for Illumina sequencing for the genome assembly. We generated this population by collecting a set of JT4 monogenic hermaphrodites and raising them in the laboratory for 6 generations (Weeks 2004). Because monogenic hermaphrodites cannot interbreed and can only produce hermaphroditic offspring, the resulting population was the exclusive product of selfing for 6 generations. Although diversity may exist between individuals in this population, each individual is highly homozygous. We sampled a single hermaphrodite from this population and expanded it to obtain the isohermaphrodite line (JT4(4)5-L) and used the line for sequencing.

**Figure 1:**
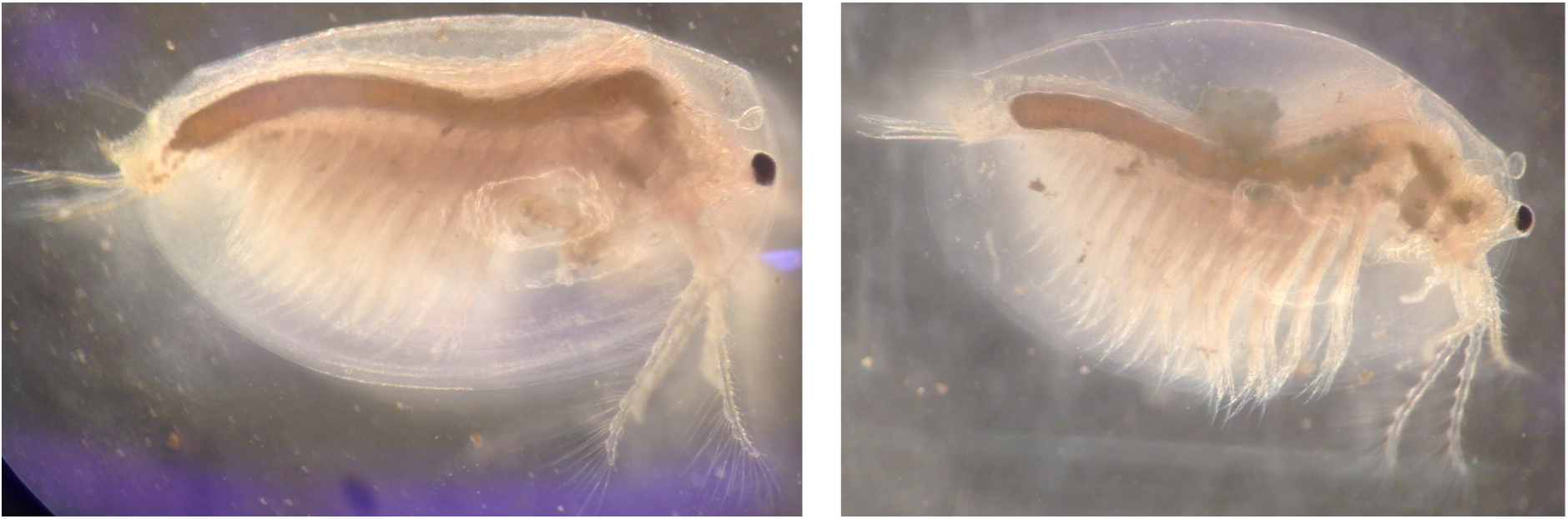
A male clam shrimp (left), and a hermaphrodite clam shrimp (right). Both are exemplars of the E. texana species. Note the presence of clasping arms on the male –these are required for nonself-fertilized sex, and the presence of a brood pouch along the dorsal surface of the hermaphrodite.

### Inbred shrimp populations sampled for genome assembly

We generated the inbred, isohermaphrodite shrimp population JT4(4)5-L from the inbred JT4(4)5-L population generated by Weeks 2004. The JT4(4)5-L population has been inbred in the laboratory (full selfing) for 6 generations, and was used for all gDNA sequencing.

### Library preparation and sequencing

#### Illumina library for genome assembly

DNA for Illumina sequencing was extracted from 50 inbred monogenic hermaphrodites from the JT4(4)5-L strain. We performed the Illumina Truseq library preparation protocol. We chose this method over Nextera library preparation for the library for genome assembly for two reasons: first, Nextera library preparation has been shown to produce a bias in coverage that can cause problems during genome assembly (Lan et al. 2015); second, the Covaris shearing used in the Truseq protocol allowed us to control the fragment length of the DNA to produce pseudo long reads obtained by joining overlapping read pairs (we refer to these a ‘pontigs’ for paired-contigs). In order to produce an average pontig fragment length of 150bp, we used the following Covaris shearing settings: 60 seconds × 6 at 10% duty cycle, 5 intensity, 200 cycles per burst. We size selected the final library on an agarose gel to get the desired 150bp read length. We ran one lane of paired-end 100bp Illumina sequencing on an Illumina HiSeq 2500, producing 124.9Gb of sequence data.

#### PacBio library for genome assembly

We followed the general protocol outlined in (Chakraborty et al. 2016) to generate the PacBio library used here. We homogenized 265 inbred monozygotic hermaphrodites from the JT4(4)5-L strain in liquid nitrogen using a mortar and pestle. We then extracted DNA using the Qiagen Blood and Cell culture DNA Midi Kit (Qiagen, Valencia, CA, USA). We made two modifications to the protocol: first, we incubated the tissue powder in the mixture of G2 buffer, RNaseA, and protease for 18 hours, rather than the 2 hours listed in the protocol; second, we doubled the RNaseA added from 19ul up to 38ul, and halved the protease added from 500ul to 250ul. We made these changes based on the presence of RNA in earlier attempts to use this kit. After gDNA extraction, we sheared the gDNA using a 1.5-inch, 24-gauge blunt tipped needle for 20 strokes. We visualized both the original gDNA and the sheared DNA using field inversion gel electrophoresis as in Chakraborty et al. 2016. We size selected the DNA using a 15kb-50kb cutoff using the BluePippin gel electrophoresis platform (Sage Science, Beverly, MA). We prepared the sequencing library using 5ug of this product, and then size selected again using a 15kb-50kb cutoff on the BluePippin gel electrophoresis platform. This produced a total of 0.149 nmol of library. We sequenced this library using 10 SMRTcells on the PacBio RS II sequencer, producing 6.7Gb of sequence data and a read length N50 of 15.2kb.

#### RNA sequencing

Individuals for RNA sequencing were derived from the WAL wild population. Adult males and hermaphrodites were sequenced separately. RNA extraction was performed using Trizol (Chomczynski and Sacchi 1987). We cleaned the RNA using RNeasy Mini columns (74104, Qiagen) following the manufacturer’s protocols, and then used this RNA to generate Illumina TruSeq RNAseq libraries according to the standard Illumina protocol. The male and hermaphrodite libraries were sequenced using one lane each of paired end 100 bp Illumina sequencing. We generated 23Gb of sequence data for males and 23Gb of sequence data for hermaphrodites.

#### k-mer counting

We generated k-mers using Jellyfish, v. 1.1.6 (Marçais and Kingsford 2011). We counted all 25-mers in the joined, but uncorrected, pontigs, then identified a local maximum coverage of 76X, then computed the genome size using the following formula:

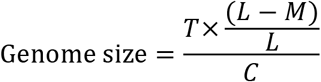

Where *T* = 15.7 *Gb* = total basepairs of pontig data, *L* = 112.7 = mean read length, *M* = 24 = mer length – 1, and *C* = 76 = coverage (cf. Lamichhaney et al. 2016). This produced a genome size estimate of 144Mb. We use this genome size estimate throughout this work.

### Genome Assembly

#### Hybrid assembly

Genome assembly was performed according to the protocol established in (Chakraborty et al. 2016). We first generated “pontigs” from the PE100 reads obtained from the 150bp insert library by assembling individual read pairs. There is some evidence (cf. read joining with a third read in Gnerre et al. 2011) that such long, contiguous, error-free reads are slightly better for genome assembly than trimmed paired reads. We generated pontigs using the *fq-join* function in *ea-utils* (Aronesty 2013), and then used *Quake* (Kelley, Schatz, and Salzberg 2010) to error correct the pontigs. We then assembled the corrected pontigs using *Platanus* (Kajitani et al. 2014), a De Bruijn graph assembler, with its default settings. This produced an assembly with an N50 of 5.2kb. We input this assembly, plus the raw PacBio reads, into *DBG2OLC* (Ye et al. 2016). The input dataset producing the highest contiguous assembly was identified via a set of hybrid assemblies using a range of quality cutoffs – we tested every whole numbered quality cutoff from 82% to 92%, and, in keeping with (Chakraborty et al. 2016), downsampled each PacBio dataset down to the longest 30X. The 85% cutoff produced the highest N50 of 1.92Mb and an assembly size of 120Mb. All N50s are summarized in supplementary table 1.

#### PacBio-only assembly

We used *Celera* 8.2, release candidate 3 (Myers et al. 2000), to generate the PacBio-only assembly, using the specfile listed in the supplementary materials (supplementary text). The assembly had an N50 of 3.4Mb, and a genome size of 126Mb.

#### Assembly merging

We used *Quiver* (Chin et al. 2013) to correct both the hybrid assembly and the PacBio assembly, then performed merging using *quickmerge* (Chakraborty et al. 2016). We used the following command line settings:

~~~
python merge_wrapper.py-pre
merged_quivered_shrimp_assemblies-hco
5.0-c 1.5 /path/to/quivered/hybrid
/path/to/quivered/pbonly
~~~

Here, -hco refers to the stringency with which seed high confidence overlaps are filtered, and -c refers to the stringency with which other HCOs are filtered. After merging, we corrected the resultant assembly by using *Quiver* again. In keeping with the *Quiver* standard practices, we ran *Quiver* on this assembly one more time, and then quantified differences between the assemblies using *MUMmer* (Kurtz et al. 2004). We noted a decrease in the number of SNPs and indels identified between the final two *Quiver* runs, so we took the final quivered assembly as our final assembly.

### Annotation

We used *Trinity* (Grabherr et al. 2011) and *Augustus* (Stanke and Waack 2003) to generate an annotation of the genome assembly. We ran *Trinity* three times: once for the male RNAseq data, once for the hermaphrodite RNAseq data, and once for the combination of both males and hermaphrodites. We used a custom script to convert *Augustus* data into a generic gff3 file, and another custom script to identify 4-fold degenerate sites based on the same annotation. We used *BLAST* (Altschul et al. 1990) to align the entire *Drosophila melanogaster* proteome against the *Augustus*-generated shrimp CDS and vice-versa. Mutual best hits with an e-value below 10^−5^ were considered significant. We tentatively assert that these genes are correctly annotated, and that they are orthologous or paralogous to genes in *D. melanogaster*.

### Differential expression analysis

We identified differences in expression between males and hermaphrodites using *Tophat* (Trapnell, Pachter, and Salzberg 2009) and the *DESeq* 1 package (Love, Huber, and Anders 2014). *Tophat* was used for transcript counting, while *DESeq* was used for differential expression analysis. Because we did not have replicated RNAseq data, we used the ‘blind’ method to estimate dispersion using the following R code:

~~~
cds <-
estimateDispersions(cds,method=blind,sharin
gMode=c(fit-only))
~~~

We then identified differences between the base means of the ‘male’ and ‘herm’ groups using the modified binomial test featured in *DESeq*, using the following R code: res=nbinomTest(cds,herm,male)

### *BLAST* annotation

We annotated all gene functions using *blastp* to align the *E. texana* genes to the *D. melanogaster* NCBI protein database, and vice versa. We regard the mutual best hits (those pairs that had e-values below 10^−5^ in both directions, and that paired in both *BLAST* directions) as the annotations in which we were most confident. In the 13 peaks of high interest discussed below, we annotated the genes that did not have mutual best BLAST hits in *Drosophila melanogaster* by taking the most significant BLAST hit for each gene (identified using *blastp* against the *D. melanogaster nr* protein database) and assigning that putative identity to the gene of interest.

### Hox gene annotation

We identified an initial set of HOX genes using mutual best hit BLAST and found 6 apparent HOX genes spread across two contigs (C0002 and C0007, Fig. 2). We then used a protein-protein BLAST (BLASTP, cutoff = 10^−5^) of all *E. texana* annotated genes onto all *D. mel* annotated genes, and identified five more genes that BLASTed to the *D. mel* HOX region. We aligned all protein sequences with Clustal-Omega (Sievers et al. 2011, Goujon et al. 2010, McWilliam et al. 2013, default settings), and then built a tree using MEGA v. 7.0.26 (Kumar et al. 2016,). Our MEGA settings were maximum likelihood tree, using only conserved residues, 300-iteration bootstrap consensus. We called any *E. texana* gene with only one *D. mel* HOX gene in its sister clade as an ortholog of the *D. mel* HOX gene. Finally, we ran a tBLASTx of the *D. mel* HOX genes against the *E. texana* genome to identify possible unannotated HOX genes (cutoff = 10^−5^). We identified *Scr* as the ortholog of the two unannotated *E. texana* genes by aligning their genomic regions, and all *E. texana* and *D. mel* HOX CDS sequences in Clustal-Omega, then calling them orthologs using the same criterion as above.

**Figure 2:**
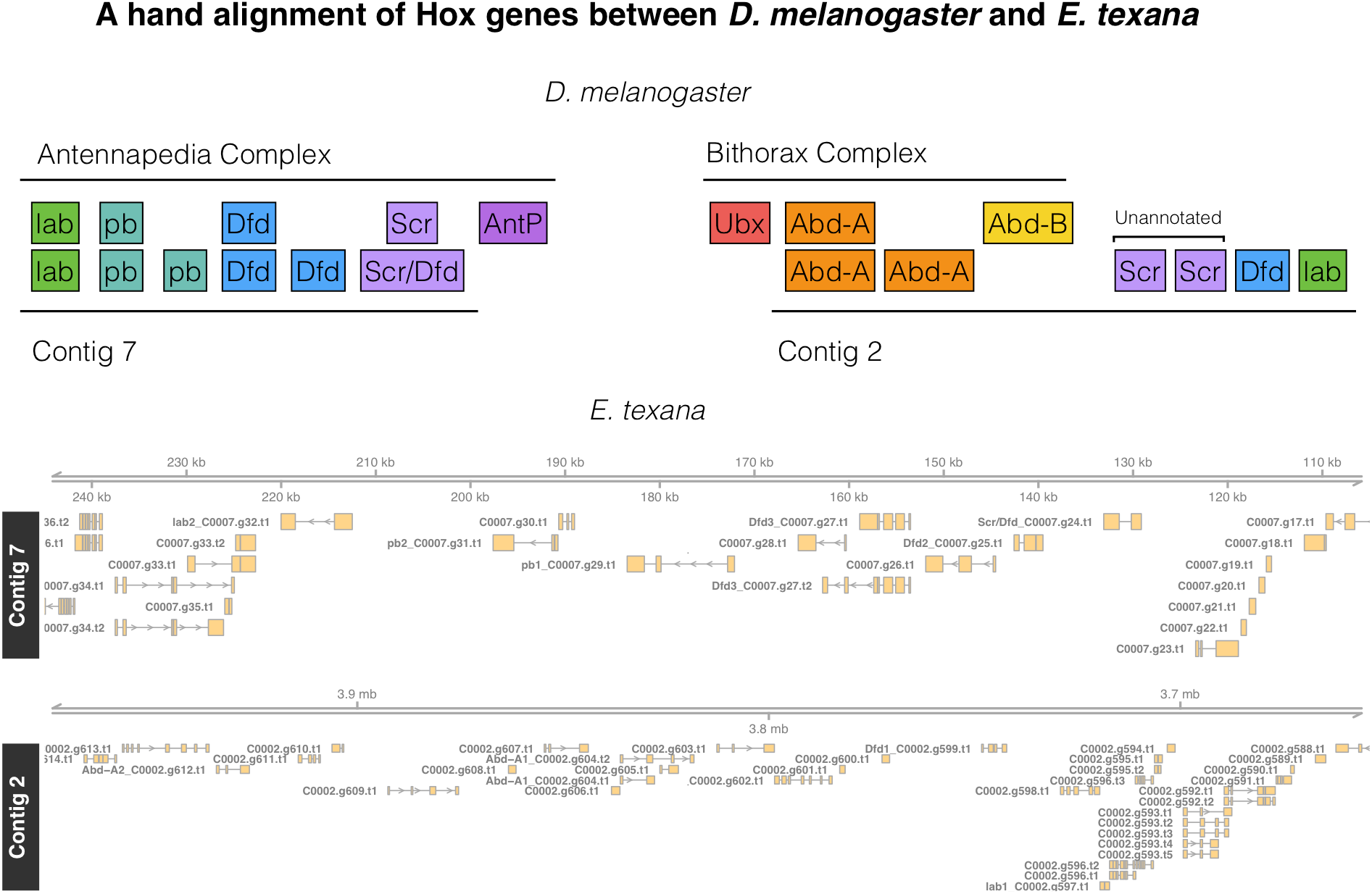
Top: The D. melanogaster HOX regions hand-aligned against the E. texana HOX regions. The ortholog identities of the E. texana HOX genes are established via bootstrap consensus maximum likelihood trees in MEGA. Note the similarity between the Antennapedia complex and Contig 7, and note that Contig 2 appears to be a combination of a copy of the Antennapedia complex and a portion of the Bithorax complex. Bottom: a visualization of the genome regions identified above. In this bottom panel, genes have been renamed for clarity. Genes that correspond to a hox gene are renamed in the figure as “DrosophilaName_E.texana name” with the Drosophila gene name prefixed to the E. texana gene name. Each instance of a given Drosophila name is numbered. To extract the correspond gene from E texana annotation files the Drosophila prefix should be removed.

**Figure 3:**
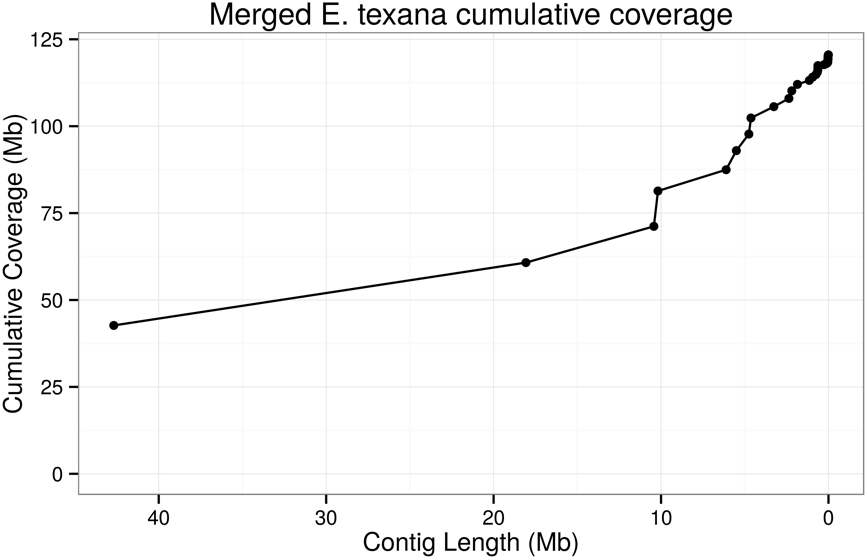
A plot of cumulative genome coverage of the E. texana genome assembly by contig. As the plot progresses from left to right, the contig lengths are added to the cumulative coverage in order from largest to smallest. A high quality assembly should achieve a high cumulative coverage with a small number of contigs. Here ~80% of the assembly is contained in contigs larger than ~5Mb.

## Results

### Genome assembly

We assembled the genome using both the hybrid approach suggested by DBG2OLC (Ye et al. 2016) and the PacBio-only approach used in *PBcR* (K. Berlin et al. 2015), and then merged the two assemblies using *quickmerge* (Chakraborty et al. 2016) to produce the final assembly. The genome assembled into 112 contigs totaling 120Mb. These contigs had an N50 of 18Mb. A plot of cumulative coverage versus contig length () demonstrates that a substantial portion (85%) of the genome is contained in the eight largest contigs. The largest contig is 41Mb in length. This level of contiguity is a dramatic improvement for vernal pool research: the highest quality vernal pool species currently assembled is *Daphnia pulex*, with a genome size of 153Mb and a scaffold N50 of 494kb (Ye et al. 2017). Other major invertebrate genomes include the honey bee (*Apis mellifera*, contig N50 = 46kb, scaffold N50 = 997kb, Elsik et al. 2014), the *Tribolium* beetle (contig N50 = 41kb, scaffold n50 = 992kb, Richards et al. 2008), and the argentine ant (*Linepithena humile*, contig N50 = 35kb, scaffold N50 = 1.3Mb, Smith et al. 2011). Note that scaffold N50 differs from contig N50 in that scaffolds are inferred by joining contigs with gaps, while contigs are gapless; thus, the difference between the assemblies is more dramatic than the numbers seem to indicate.

The observation that the estimated genome size is 144Mb, and the final assembly size is 120Mb, indicates that some portions of the genome were not assembled. This is ordinary in genome assembly, as highly repetitive heterochromatin regions tend to be impossible to assemble with current technology. For instance, the *Drosophila melanogaster* genome is estimated to be 175Mb in size (Ellis et al. 2014), yet the *D. melanogaster* assembled genome (easily among the best higher eukaryote assemblies) is “only” 143Mb (Santos et al. 2015).

Two lines of evidence lead us to have confidence in this genome assembly: the quality of other genome assemblies produced using similar data and the same bioinformatics pipeline, and empirical evidence of the quality of this assembly. The genome assembly pipeline used in Chakraborty 2016 (Chakraborty et al. 2016) has been thoroughly evaluated under a variety of genome size and coverage circumstances, and the genome size and coverage of these test assemblies match very closely to the genome size and coverage of our *E. texana* assembly. In particular, the Chakraborty 2016 assembly that used 39X of coverage to assemble a 140Mb genome had an assembly N50 of 6.69Mb, only 3194 misassemblies, and 12.25 mismatched bases per 100kb. Empirical evidence of the quality of a never-before-assembled genome is difficult to acquire, but we can report on the fraction of the Trinity-assembled (Grabherr et al. 2011; detailed below) RNAseq-derived transcripts that are present within the final assembly. We find that, if we use transcripts assembled entirely from RNA from hermaphrodites of the reference strain JT4(4)5-L, 98.9% of the transcripts align with above 92% identity, according to *BLAT* (Kent 2002). Interestingly, using the entire RNAseq dataset, which contained both the hermaphrodites from the reference strain and males from the WAL strain, produced 95.5% successful alignment, which opens the possibility that some genes are present only in some male fraction of the genome not sampled in our WW hermaphrodite. Unfortunately, this difference could alternatively be strain-specific, rather than male-specific, with no simple way to differentiate those possibilities without further experimentation.

*Repeatmasker* identified 624 SINEs, 16,044 LINEs, 2302 LTRs, 24817 DNA elements, and 88928 unclassified elements, together making up 26.4% of the genome. This contrasts with the relatively low rate of repetitive elements in *D. melanogaster*, at 3.9% (Kaminker et al. 2002). That said, a large portion of this repetitive sequence is ‘unclassified’; if we remove the unclassified repeats from the count, only 9.8% of the genome consists of interspersed repeats. Other (non-interspersed) repeats make up 5.1% of the genome.

### Annotation and differential expression

We collected one lane of Illumina RNAseq data from 25 male clam shrimp from the WAL wild population, and another lane from 25 inbred monozygotic females from the JT4(4)5-L population (the reference population used for the assembly). We used a combination of *Trinity* (Grabherr et al. 2011) and *Augustus* (Stanke and Waack 2003) to generate an annotation. We did three runs of *Trinity* – one run using only the males, one run using only the hermaphrodites, and one run using both together. The combined run produced 85,721 transcripts, while the male and hermaphrodite runs produced 77,257 and 55,845 transcripts, respectively. We ran *Augustus* using the combined run to generate gene predictions for *E. texana*. This generated a total of 17,667 genes and 23,965 transcripts. Of these genes, 5,438 were found to be mutual best hits with known *D. melanogaster* genes.

In order to validate our annotation and assembly, we attempted to identify HOX genes in the clam shrimp genome, and compare their order to that of the HOX genes in *D. melanogaster*. HOX genes are an interesting test case as they are important in development, they are believed to cluster in two different chromosomal regions in invertebrates, their order tends to be conserved across all animals, and that order reflects where they are expressed along the anterior/posterior axis (reviewed in Duboule 2007). We identified HOX genes using mutual best hit BLAST and found 6 apparent HOX genes spread across two contigs (C0002 and C0007, Fig. 2). We then used a protein-protein BLAST of all *E. texana* annotated genes onto all *D. mel* annotated genes, and identified five more genes that BLASTed to the *D. mel* HOX region. We removed one of these genes (C0002.g600) from the analysis because, upon multiple alignment with Clustal-Omega (Sievers et al. 2011, Goujon et al. 2010, McWilliam et al. 2013), there was no evidence that it contained a HOX motif. Finally, we ran a tBLASTx of the *D. mel* HOX genes against the *E. texana* genome to identify possible unannotated HOX genes in the region, and found two more candidates. We took this collection of 12 genes, found orthologs between *E. texana* and *D. mel* using Clustal-Omega and MEGA v. 7.0.26 (Sup. Figs. 1 and 2; see also Kumar et al. 2016), and hand-ordered them relative to *D. mel*. The identity of these genes is not certain, but from our results, it appears that nearly all genes are grouped spatially with their orthologs, and the rough order of the orthologous gene groups is conserved between *D. mel* and *E. texana*, especially when comparing the *D. mel* Antennapedia complex to the *E. texana* genome (of the *D. mel* Bithorax complex, only Abd-A orthologs were identified in *E. texana)* (Fig. 2). Additionally, Contig 2 appears to contain a partial duplication of the Antennapedia locus from *D. mel*.

We next compared the RNAseq data from males and hermaphrodites to identify differentially expressed genes. We found 486 differentially expressed genes (Benjamini-Hochberg-Yekutieli (Benjamini and Yekutieli 2001) adjusted p-value below 0.05) (Fig. 4) out of the 17,667 genes identified by *Augustus*. Forty of these genes are amongst the genes with *D. melanogaster* orthologs. Gene ontology enrichment analysis with *GOrilla* (Eden et al. 2009) indicates an enrichment of the following GO terms based on the rank order of significance of differential expression (GO terms with a Benjamini-Hochberg corrected *p*-value below 0.05 are listed): structural constituent of cuticle, chitin binding, structural constituent of chitin-based larval cuticle, structural constituent of chitin-based cuticle, carboxypeptidase activity, chitin deacetylase activity, and association with the condensin complex, extracellular region, and DNA packaging complex (Sup. Table 2). Hermaphrodites have both testes and ovaries, while males have only testes; additionally, hermaphrodites typically store up to several hundred large eggs in their carapace prior to ovipositioning (Weeks, Marcus, and Alvarez 1997). These two large phenotypic differences between males and females are likely to drive many of the observed expression differences.

### Sex locus localization

The quality of the clam shrimp genome assembly allows us to identify the contig harboring the sex-determining locus of *E. texana*. Previous analyses of allozymes and microsatellites (S C Weeks 2004; S. C. Weeks et al. 2010) indicate the sex determining locus is linked to several markers, with at least three markers so tightly linked that they can be used to genotype the sex locus status of individuals (ZZ vs. ZW vs. WW) with relatively high accuracy. We used *BLAST* (Altschul et al. 1990) to align the sequences of the four best such markers (the allozyme *Fum* and microsatellites *CS8, CS11, and CS15*) to the *E. texana* assembly (Sup. Fig. 3). We found that the three microsatellite loci aligned to our largest (41Mb; contig 1) contig, while the allozyme *Fum* aligned to a smaller 1Mb contig (that we speculate would join contig 1 in a more contiguous assembly). The order (in the assembly) of the three microsatellite markers that map to contig 1 does not agree with the order inferred genetically in (S. C. Weeks et al. 2010; see figure), indicating a problem with either the mapping or the assembly. We are relatively confident in the quality of our assembly, and there is reason to think that the mapping could be incorrect. Weeks 2010 found a very high rate of recombination between three microsatellites when looking at male meiosis. Specifically, he observed recombination distances of 94 and 73 cM; recombination fractions indistinguishable from free recombination. In contrast, in hermaphrodites, adjacent markers were separated by a very small number of recombinants with only approximately 5 total crossover events inferred in 170 individuals. We posit that recombination does not occur in amphigenic hermaphrodites, or occurs very seldom, and that much of the inference of marker order may actually be due to a low (~1%) rate of mis-genotyping of the microsatellite markers. Our highly contiguous genome allows for future experiments to determine if indeed amphigenic hermaphrodite experience recombination in *E. texana*.

**Figure 4:**
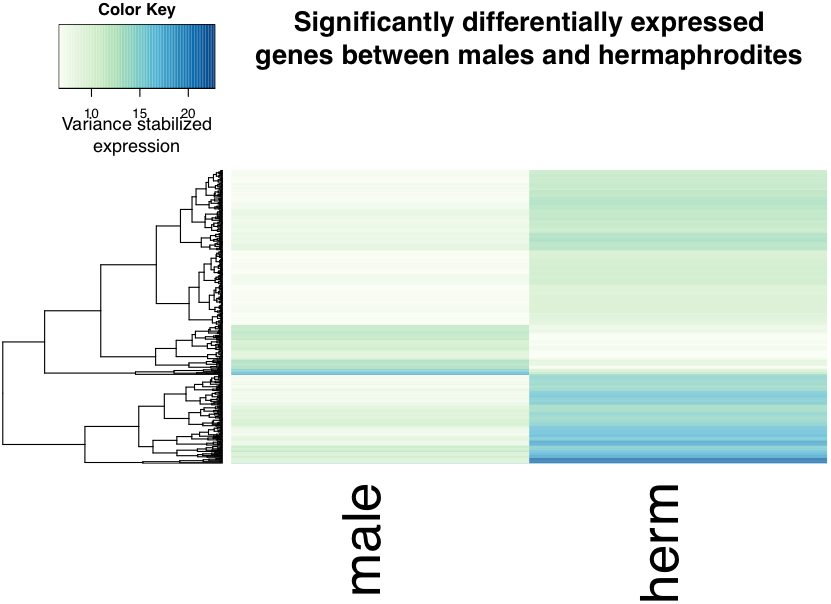
A heat map of expression for genes differentially expresses between males and hermaphrodites (adjusted *p* < 0.05). Note the small portion of genes that have nearly zero expression in males, and high expression in hermaphrodites.

It is important to note that the genome assembly was produced using data from WW hermaphrodites. Thus, the male version of the sex-determining locus is not expected to be present in the genome assembly. This may make detection of the sex-determining locus more difficult, depending on the divergence of the ‘Z’ and ‘W’ versions of the sex locus. If the two loci are highly diverged, they may not align to each other; on the other hand, if they are not highly diverged, they may align to each other, but show a signal of increased polymorphism. A future *de novo* assembly of a male will help elucidate the location of the sex determination locus.

### Residual Variation and Assembly Errors

We aligned the Illumina data from the inbred JT4(4)5-L line used for the genome assembly to the reference genome and observed SNPs at a rate of 0.00018 per bp. This indicates, as expected, a very low SNP rate within the inbred strain we sequenced (Supplementary Figure 4). If JT4(4)5-L was not fully inbred then we expect runs of heterozygous sites, whereas isolated SNPs are likely assembly errors. Consistent with this prior belief there are notable differences in patterns of heterozygosity amongst the contigs. The largest three contigs are almost completely free of heterozygosity (0.000024 SNPs per bp), reflecting a very low assembly error rate at with respect to point mutations. In contrast, the fourth contig and several others generally have higher levels of heterozygosity (contig 4: 0.00021 SNPs per bp). Sixty-four percent of the heterozygosity in the genome is contained in the 26 most SNP-dense contigs, which account for only 5.6Mb of the genome. Thus, most of the genome is nearly heterozygosity free with blocks of residual heterozygosity. We speculate that these small contigs with high levels of heterozygosity could be mis-assembled, leading to incorrect read mapping that appears as heterozygosity, or regions that did not become homozygous following inbreeding that then failed to assemble adequately because of the heterozygosity therein.

## Discussion

### On non-model organisms and genome assembly

One of the long standing assumptions in genomics is that high quality whole-genome genetic analysis is not possible with nonmodel organisms because of the lack of genetics resources available for such systems, such as genome assemblies and annotations. Before the advent of high-throughput sequencing (i.e., Illumina sequencing), non-model genome assembly was prohibitively expensive. The human genome project cost approximately $3 billion, while the Celera human genome assembly was seen as comparatively affordable at $300 million. The advent of Illumina sequencing and De Bruijn graph assembly dropped the cost of genome assembly to on the order of $10,000 - depending on the genome size and complexity - but the contiguity of these assemblies tended to be low because of the short length of Illumina-type reads. Thus, most arthropods, with the exception of *D. melanogaster*, have had low contiguity genome assemblies when they have assemblies at all. One of the most studied insects, the *Heliconius melpomene* butterfly, is a representative example. Its 454 and Illumina-based assembly, published in 2012 by a large consortium, had an N50 of 277kb (Heliconius genome consortium, 2012), which was considered very respectable contiguity for a non-model assembly at that time. In 2016, PacBio sequencing and linkage analysis was used to bring the N50 of *H. melpomene* to 2.1Mb, highlighting the advances possible with long read sequencing technology (Davey et al. 2016). Still, outside of the insects, high quality genome assemblies are rare. *Daphnia pulex*, which has been used as a model organism for many years, has an assembly with a scaffold N50 of 470kb (Colbourne, Singan, and Gilbert 2005). We have now generated what is, to our knowledge, the most contiguous crustacean assembly ever completed. Here, we demonstrate that the generation of a genome for a new model organism is not necessarily difficult or costly. Modern sequencing techniques (i.e., PacBio) allow for de novo genome assembly of a ~200Mb genome for ~$10K USD. A preliminary genome annotation using RNAseq for a handful of tissues can be accomplished for ~$3K USD. This combination of factors makes genomics in non-model systems an attractive target for evolutionary biologists.

We present here a *de novo* whole genome assembly for *E. texana* with an N50 of 18Mb. This genome will be a useful resource for the vernal pool research community, and will elevate the status of clam shrimp as an emerging model organism. Additionally, we present a draft annotation of the genome that allows for accurate identification of genic, intergenic, etc. regions, as well as homology-based comparisons with genes in other species. Finally, we carried out an initial analysis of differential gene expression between males and hermaphrodites and identify some gene ontology terms that seem to be associated with differential expression between males and hermaphrodites.

### The proto-sex chromosome

Much effort has gone into identifying the structure of the sex locus in individuals with recently derived sex chromosomes (Zhou and Bachtrog 2012, Charlesworth 2012). *E. texana* is androdioecious, but is believed to be descended from a dioecious ancestor that was ancestral to the entire *Eulimnadia* clade (S. C. Weeks et al. 2009). Linkage analysis has indicated that the sex-determining region is likely to be a large autosomal linkage group or a “proto-sex” chromosome. We identified a single contig that contained all but one of the previously identified sex-linked markers. This contig likely harbors the sex determining linkage group. Linked genetic markers were spread across the entire 42-Mb contig, and the order of the markers differed from the order predicted by linkage mapping. It is not clearly relevant to the evolution of sex chromosomes, but it is an interesting observation that the sex chromosome represents roughly a third of the clam shrimp genome. We were unable to identify the sex-determining locus within this chromosome, since it is possible that hermaphrodites do not recombine, as is the case in *Drosophila melanogaster* (Lenormand 2003) and other organisms. A lack of recombination in hermaphrodites would make linkage-mapping the sex-determining locus impossible. Our genome assembly should allow for new experiments using SNP markers to confirm or refute the existence of recombination in hermaphrodites and perhaps map the sex-determining locus. Alternatively, a second male specific assembly, in concert with GWAS-type approaches, may allow the sex determining region to be identified. Additionally, we cannot rule out the possibility that the entire chromosome, rather than a narrow locus is involved in sex determination.

We mapped RNAseq derived transcripts from hermaphrodites and males back to the genome assembly. Despite our ability to map ~99% of hermaphrodite transcripts back to the reference genome, ~4% of the male transcripts failed to map. Thus, there are transcripts present in males that are too distinct to map to the hermaphrodite derived genome assembly. This suggests one of three possibilities: first, there may be a genomic region that only occurs in males, which is absent from our current assembly; second, there is a region present in both male and hermaphrodite versions of the genome, but that the male and hermaphrodite alleles are too diverged from one another for male derived transcripts to map back to hermaphrodite alleles; or third, some male transcripts do not map back to the reference simply due to polymorphism segregating in this species. We note that the RNAseq data were obtained from two different strains, with the hermaphrodite strain being the same one from which the assembly is derived. A further study could elucidate which of these hypotheses is correct by generating a whole genome assembly of a male genome (or, although less informative, aligning hermaphrodite specific transcripts from the same strain the male transcripts were from back to the reference genome).

## Conclusions

We generated a highly contiguous, annotated genome assembly with an N50 of 18Mb for the clam shrimp *Eulimnadia texana*. This genome assembly allowed us to identify numerous genes with homology to genes in *Drosophila melanogaster*, and we identified a subset of these genes as being differentially expressed between males and females.

## Data deposition

All sequencing data is available at the NCBI All data will be made available at the NCBI Sequencing Read Archive and NCBI Genbank under the Bioproject “PRJNA352082”. The genome, also under this Bioproject, has the accession number “NKDA00000000”. Additional files are available at the following URL: http://www.wfitch.bio.uci.edu/~tdlong/PapersRawData/BaldwinShrimp.tar.gz. Additionally, all scripts used for analysis will be made available at the following GitHub page: https://github.com/jgbaldwinbrown/jgbutils

## Specific Acknowledgements

Thanks to the UC Irvine genomics core facility and the University of Kansas genomics facility, as well as Stuart Macdonald, for assistance in library preparation and sequencing.

